# Direct coevolutionary couplings reflect biophysical residue interactions in proteins

**DOI:** 10.1101/061390

**Authors:** Alice Coucke, Guido Uguzzoni, Francesco Oteri, Simona Cocco, Remi Monasson, Martin Weigt

**Affiliations:** Laboratoire de Physique Théorique, Ecole Normale Supérieure and CNRS-UMR8549, PSL Research University, 24 Rue Lhomond, 75005 Paris, France; Sorbonne Universités, UPMC, Institut de Biologie Paris-Seine, CNRS, Laboratoire de Biologie Computationnelle et Quantitative UMR 7238, 75006 Paris, France; Laboratoire de Physique Statistique, Ecole Normale Supérieure and CNRS-UMR8550, PSL Research University, Sorbonne Universités UPMC, 24 Rue Lhomond, 75005 Paris, France

## Abstract

Coevolution of residues in contact imposes strong statistical constraints on the sequence variability between homologous proteins. Direct-Coupling Analysis (DCA), a global statistical inference method, successfully models this variability across homologous protein families to infer structural information about proteins. For each residue pair, DCA infers 21×21 matrices describing the coevolutionary coupling for each pair of amino acids (or gaps). To achieve the residue-residue contact prediction, these matrices are mapped onto simple scalar parameters; the full information they contain gets lost. Here, we perform a detailed spectral analysis of the coupling matrices resulting from 70 protein families, to show that they contain quantitative information about the physico-chemical properties of amino-acid interactions. Results for protein families are corroborated by the analysis of synthetic data from lattice-protein models, which emphasizes the critical effect of sampling quality and regularization on the biochemical features of the statistical coupling matrices.

## I. INTRODUCTION

Across evolution, the structure and function of homologous proteins are remarkably conserved. As a consequence, neighboring residues in the threedimensional structure tend to coevolve, leading to strong constraints on the sequence variability. Direct Coupling Analysis (DCA)^1,2^, a global inference method based on the maximum-entropy principle^3,4^, successfully exploits pairwise correlations in amino-acid occurrence, which are easily observable in large multiple-sequence alignments, to infer spatial residue-residue contacts within the tertiary protein structure. This approach uses a global statistical model *P*(*a*_1_,…,*a*_*L*_) for an amino-acid sequence (*a*_1_,…,*a*_*L*_) of length *L*, whose parameters are fields/biases {*h*_*i*_(*a*)} and statistical couplings {*J*_*ij*_(*a*,*b*)}, where *a*, *b* are amino acids or alignment gaps (denoted for simplicity by {1, …, 21} throughout the paper). These parameters are learnt from site-specific amino-acid frequencies, and from the covariance between amino-acid pairs estimated from multiple-sequence alignments (MSA), which are readily available thanks to rapidly increasing sequence databases^5,6^. Contact prediction is performed by measuring the total coupling strength between two residues. The coupling matrices - inferred at high computational cost - are mapped onto simple scalar parameters, and the full information they potentially contain gets lost.

The aim of our work is to provide a better quantitative understanding of these inferred couplings. Earlier works have shown that the coevolutionary couplings derived by DCA contain an electrostatic signal^7^. In the present study, we go considerably further and show that the coevolutionary couplings also contain quantitative and interpretable biological information related to all the physico-chemical properties of amino-acid interactions, not only electrostaticity, but also hydrophobicity/hydrophilicity, Cysteine-Cysteine bonds, Histidine-Histidine and steric interactions. These interactions are consistent with knowledge-based amino-acid potentials inferred from known protein structures, such as the statistical potential derived by Miyazawa and Jernigan^8^.

To carry out our study, we first consider a set of 70 Pfam^6^ protein families from which we infer the coupling matrices. After selecting the top ranked residue pairs for each family, we analyze the mean coupling matrix and its spectral modes. Considering structural classifications and solvent exposure helps unveiling the full biological content of the coupling matrices {*J*_*ij*_(*a*, *b*)}_*a*, *b*∈{1,…,21}_. Our analysis also shows that the distribution of contact distances in the tertiary structure greatly depends on the type of interaction associated to the contact.

In a second part of the article, to better understand the effect of sampling and regularization on the previous findings, we focus on lattice proteins^9^, an exactly solvable model of proteins folding on a 3D lattice. Lattice protein indeed provide an interesting framework for testing statistical modeling approaches like DCA in a relatively realistic, and fully controllable context^10^.

## II. REVIEW OF DIRECT-COUPLING ANALYSIS OF RESIDUE COEVOLUTION

### A. Maximum-Entropy approach

A global probabilistic model *P*(*a*_1_, …,*a*_*L*_) assigns a probability to any amino-acid sequence ***A*** = (*a*_1_,…,*a*_*L*_) based on empirical frequency counts in the MSA. More precisely, in order to be coherent with the MSA, the probabilistic model is chosen to reproduce the empirical one-and two-residue amino-acid frequency counts:

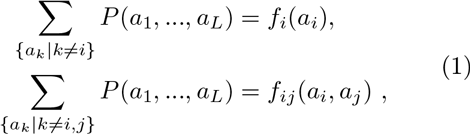

where *f*_*i*_(*a*) denotes the fraction of proteins having amino acid *a* in column *i* of the MSA, and *f*_*ij*_(*a*, *b*) counts the fraction of proteins with amino acid *a* in column *i* and amino acid *b* in column *j*. The least constrained or Maximum-Entropy (MaxEnt)^3,4^ model reproducing these observations is a Potts model with *q* = 21 (20 possible amino acids + 1 alignment gap ’-’) states, or equivalently a Markov random field:

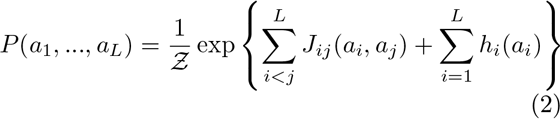

where 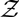 is a normalization constant (known as partition function in the context of the Potts model), and *h*_*i*_(*a*) represent site-specific local biases. Parameters {*J*_*ij*_(*a*, *b*)}_*a*,*b*=1…*q*_ are direct statistical couplings between residues *i* and *j*, taking the form of 21 × 21 matrices.

The numerical values of *J*_*ij*_(*a*, *b*) and *h*_*i*_(*a*) have to be determined such that Eqs. (1) are satisfied-leading to the approach known as *Direct Coupling Analysis* (DCA)^1,2^. From a computational point of view, it is not feasible to solve Eqs. (1) exactly: the calculations of the normalization 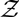 and of the marginals require to sum over all *q*^*L*^ possible amino-acid sequences of length *L*. With *q* = 21 and typical protein lengths of *L* ≃ 50 – 500, there are 10^65^ – 10^650^ possible configurations.

Several methods may be used to approximate the parameters, the computationally most efficient of which is is the mean-field approximation^2^, where the coupling matrix is the inverse of the correlation matrix. This method is closely related to the Gaussian modeling scheme used by PsiCov^11^. A more involved approximation, called *Pseudo-Likelihood Maximization* (plmDCA^12^ and GREMLIN^13,14^), is shown to outperform mean-field DCA on biological sequence data. The asymmetric version^15^ of plmDCA will be used in the present article, cf. Appendix A.

### B. Regularization and reweighting

Protein sequences are not independently and identically distributed (i.i.d.); they form a finite and usually small-size sample. Indeed, a Potts model describing a protein family with sequences of 50 – 500 amino acids requires ca. 10^6^ to 10^8^ parameters. Few protein families are large enough to directly determine these parameters, and regularization is essential to avoid overfitting. Moreover, adding a regularization term helps the hill-climbing optimization in plmDCA to rapidly find the maximum of the pseudo-likelihood. Different regularization schemes and their effects have been extensively addressed in the literature^16^.

A prior probability distribution (typically Gaussian) is considered for the model parameters, which discounts large values resulting from insufficient statistics in the original MSA. The following *l*_2_-penalty is therefore added to the log-likelihood of the data:

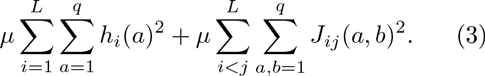

For plmDCA, the standard value of the regularization parameter is *μ* = 10^−2^ as it gives optimal results for contact prediction^12^.

On the other hand, there are strong sampling biases due to phylogenetic relations between sequenced species. This problem has been the object of previous studies^17,18^, but a simple sampling correction can be implemented by counting sequences with more than 80% identity and reweighting them in the frequency counts^2^. The number of non-redundant sequences is measured as the effective sequence number *M*_eff_ after reweighting. As a rule of thumb *M*_eff_ has to be at least 300 to enable plmDCA to predict residue-residue contacts in real proteins.

### C. Reparametrization (gauge) invariance and zero-sum gauge

The *Lq* single-residue (*f*_i_(*a*)) and 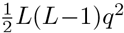 two-residue frequencies (*f*_*ij*_(*a*, *b*), *i* < *j*)^2^ estimated from the data are not independent. The former sum up to 1, and the latter have the single-residue frequencies as marginals. Therefore not all constrains in Eq. (1) are independent: The total number of non redundant parameters is actually 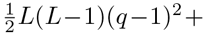 *L*(*q*−1). This number is smaller than the total number 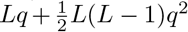 of Potts parameters *h*_*i*_ (a) and *J*_ij_ (*a*, *b*) appearing in Eq. (2). The model is therefore over-parametrized, a fact referred to as gauge invariance in physics language. We can reparametrize the model without changing probabilities using an arbitrary *K*_*ij*_(a), 1 ≤ *i*,*j* ≤ *L*,*a* ∈ {1, …, 21}:

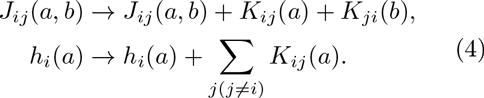

The inferred fields and couplings will be expressed throughout this paper in the so-called “zero-sum gauge”, in which 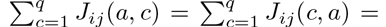 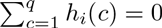 for all amino acid *a* and all positions *i*, *j*. In practice, the couplings *J*_*ij*_(*a*, *b*) can be simply put in the zero-sum gauge through

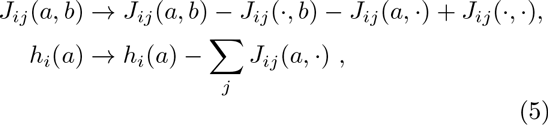

where *g*(·) denotes the uniform average of *g*(*a*) over all 21 amino acids + gap symbols *a* at fixed position. The zero-sum gauge minimizes the Frobenius norm of the coupling matrices, which is used as a scalar measure of the coupling strength. It allows for the ranking of residue pairs (*i*, *j*) in order to predict residue-residue contacts^1,12,19^.

### D. Contact prediction

After having estimated the parameter values of the DCA model *P*(*a*_1_, …, *a*_*L*_), each residue pair (*i*,*j*) is characterized by a 21 × 21 matrix {*J*_ij_(*a*, *b*)}_*a*,*b*∈{1,…,21}_. To measure the coupling strength of two sites, the inferred {*J*_*ij*_(*a*, *b*)} has to be mapped onto a scalar parameter. These parameters will then be ranked to perform a contact prediction: The bigger they are, the higher is also the probability that *i* and *j* are in contact in the model. It has been observed that a modified score - the Frobenius norm *F*_*ij*_ of the coupling matrix adjusted by an *Average Product Correction* (APC) term - improves contact prediction^12^:

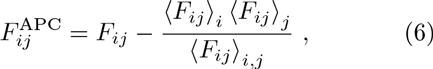

where the mean ⟨.⟩ denotes positional average over single (*i*) or double (*i*,*j*) sites. To compute this score, the couplings are first shifted to the zero-sum gauge described in Eq. (5) after the inference by plmDCA.

### E. The Miyazawa-Jernigan statistical potential

**FIG. 1:**
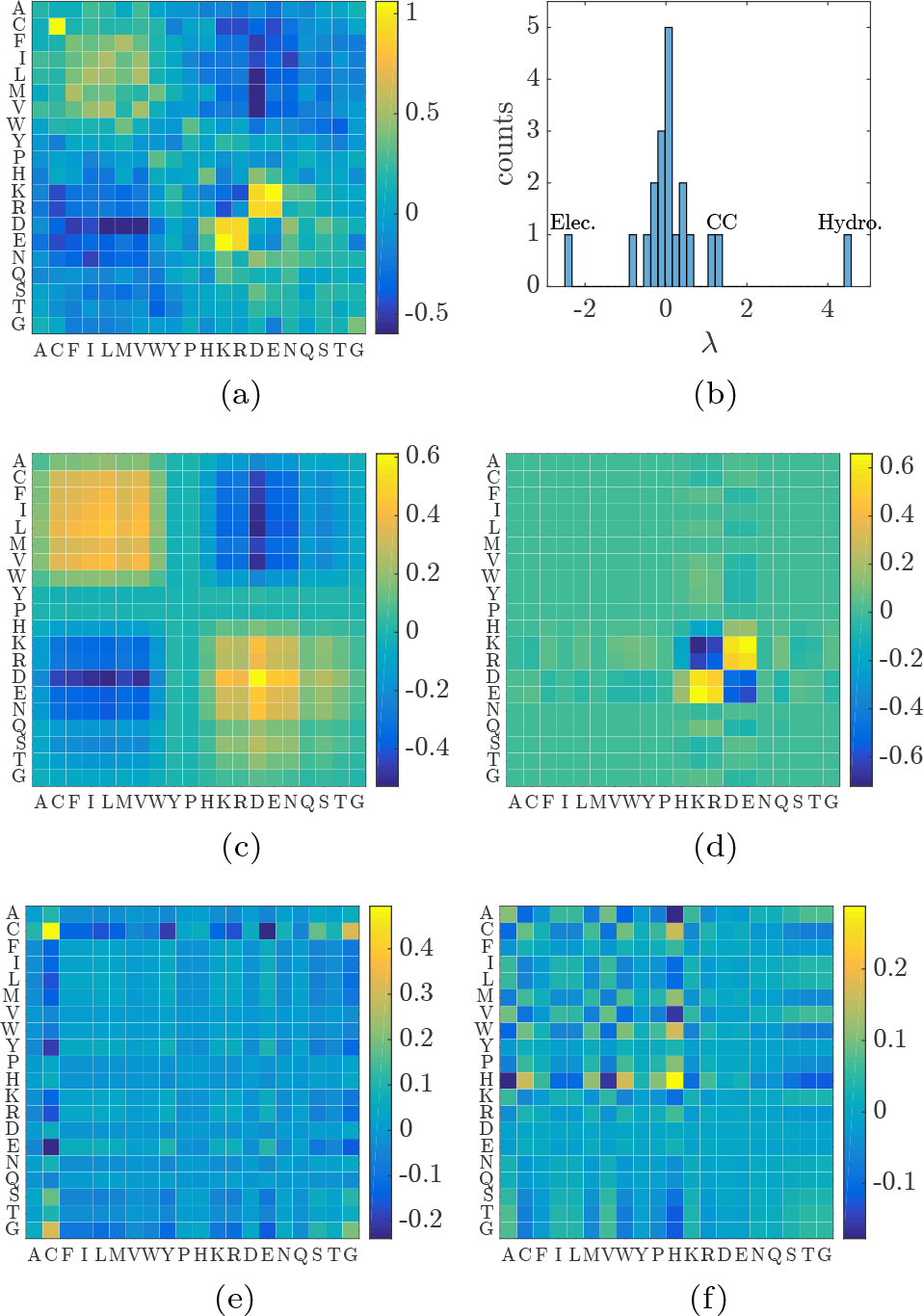
(a) Miyazawa-Jernigan (MJ) energy matrix 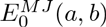. (b) Spectrum of the MJ matrix. MJ’s 3 largest spectral modes, displaying physico-chemical interactions: (c) hydrophobicity-hydrophilicity (λ^(1)^ = 4.55), (d) electrostaticity (λ^(2)^ = −3.51), (e) Cysteine-Cysteine (λ^(3)^ = 1.28), and (f) Histidine-Histidine (λ^(4)^ = 1.04) signals.

Developed from the 1980s, the Miyazawa-Jernigan (MJ) knowledge-based potential *E*^*M J*^ (*a*, *b*) was derived from the statistics of amino acids in contact in known 3D protein structures. This 20×20 interaction matrix reflects the physico-chemical properties of the amino acids, torsions angles, solvent exposure and hydrogen bonds geometry^8^. In contrast to more detailed potentials including also, e.g., the residue distance, the MJ interaction matrix is a natural starting point for comparison with the DCA-derived coupling matrices. Panel (a) of Fig. 1 displays 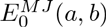, the 20 × 20 matrix provided by Miyazawa and Jernigan in 1996^20^, upon transformation into zero-sum gauge with the help of Eq. (5), to compare with DCA couplings later on. It has also been multiplied by a factor −1 to comply with the standard convention that attractive interactions are positive, and repulsive ones are negative:

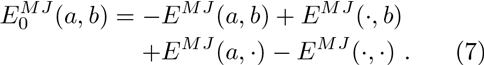

In this specific gauge, the spectrum of the MJ matrix shows a few significant eigenvalues (Fig. 1 panel (b)).

Panels (c) to (f) display the first spectral projections of the MJ matrix 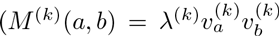, *k* = 1…4, see Eq. (10) below). They are localized on particular amino acids according to physicochemical interactions. Panel (c) is related to hydrophobicity/hydrophilicity: amino acids from A to P are hydrophobic, whereas the rest are hydrophilic. Hydrophobic amino acids tend to form contact with other hydrophobic amino acids but not with hydrophilic ones, according to the signs of the corresponding entries. Panel (d) is related to electrostaticity: amino acids K, R and H are positively charged whereas D and E are negatively charged. Panel (e) is localized on the Cysteine-Cysteine entry, as those amino acids tend to form strong chemical disulfide bounds where paired with each other. Finally, panel (f) shows the fourth spectral mode of the MJ matrix, localized on the Histidine-Histidine entry, forming like-charged contact pairs^21^.

The eigenvalues corresponding to hydrophobicity/hydrophilicity (λ^(1)^ = 4.55), the Cysteine-Cysteine (λ^(3)^ = 1.28) and Histidine-Histidine interactions (λ^(4)^ = 1.04) are positive, describing an attractive interaction between like amino acids. On the other hand, the eigenvalue corresponding to electrostaticity (λ^(2)^ = −3.51) is negative, reflecting the attraction between charges of opposite sign, and repulsion between like charges.

## III. RESULTS ON PROTEIN SEQUENCES DATA

### A. Method

We consider 70 protein families from the Pfam database^6^, containing enough sequences (*M*_eff_ > 500) to guarantee a good inference (sufficient sampling for plmDCA), and possessing at least one X-ray crystal structure of resolution below 3Å in the Protein Data Bank^22^ (PDB); the complete list can be found in Supplementary Section IV. For each Pfam family *n* we infer with the plmDCA method^15^ the 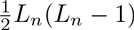 (with *L_*n*_* being the aligned length of the proteins in family *n*) coupling matrices at standard regularization (*μ* = 10^−2^), and transform them into zero-sum gauge. The top ranked pairs (*i*,*j*) of residues (according to the *F*^*APC*^score defined in Eq. (6)) are selected until a rate of 20% of false-positive contact predictions is reached within the selection. Then, only the true-positive predictions (contacts in the tertiary structure) are kept in the selection 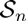. The number of selected pairs 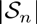 thus depends on the Pfam family *n*. We obtain the global selection of residue pairs 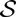 by assembling the selected pairs of each Pfam family together: 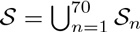 with 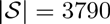.

Here, a residue pair is considered to be a true positive prediction if its minimal heavy-atom distance is below 6Å in the protein structure (the method used to define the contact map from the protein crystal structures is described in Appendix B). To avoid both trivial contacts and strong but uninformative “gap-gap” signals, we also impose a minimum separation |*j* – *i*| > 10 along the protein backbone. Indeed, gaps in the MSA are not generally modeled well by DCA methods, as they tend to come in long stretches, giving rise to artificially high couplings for closer sites on the backbone^23^.

In the following, we consider the mean matrix

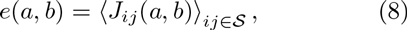

where 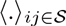 denotes the mean over all residue pairs in the above-mentioned selection 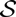, all Pfam families taken together. The matrix *e* is subsequently symmetrized, as any non-symmetric features of amino-acid interactions originate from finite-sampling effects in the selection,

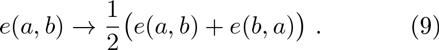

The average coupling matrix *e* is already in the zero-sum gauge, since the couplings *J*_*ij*_ are. By considering the mean matrix, we expect site specificities and finite-sampling noise to be averaged out, while the joint global interaction modes should be prominently displayed.

We define the spectral mode *k* of *e* by

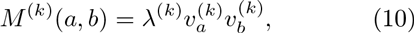

where {λ^(*k*)^,*υ*^(*k*)^}_*k* = 1…21_ are the eigenmodes of *e*, with the eigenvalues λ^(*k*)^ ranked in decreasing order in absolute value.

### B. The coupling matrices contain biologically relevant information

**FIG. 2:**
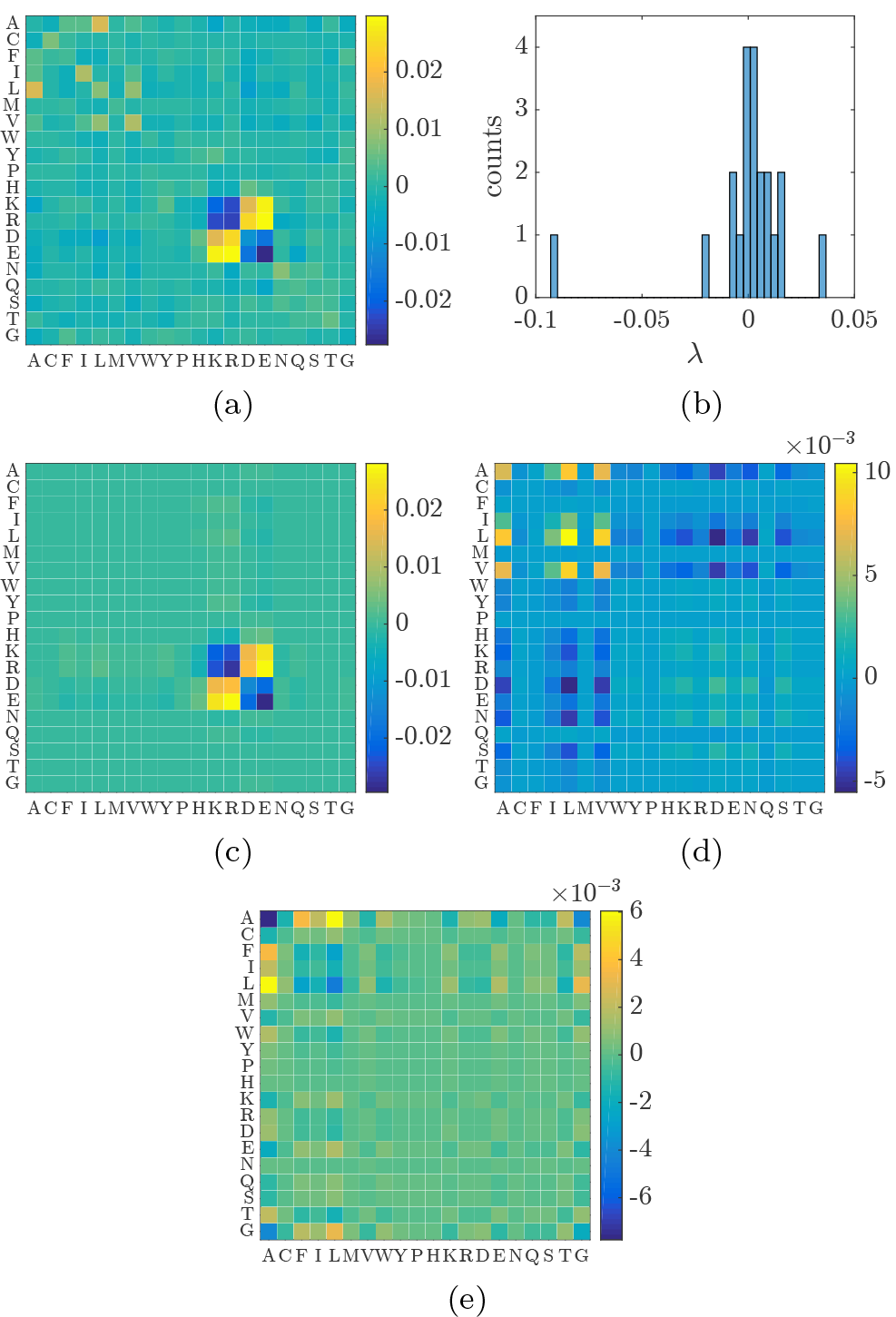
(a) mean matrix *e*(*a*, *b*) over all residue pairs in the selection, taking all Pfam families together.(b) Histogram of the spectrum of *E*, dominated by three eigenvalues. (c) First spectral mode of *E* (λ^(1)^ = −0.0923), displaying the electrostatic interaction. (d), (e) Second (λ^(2)^ = 0.0363) and third (λ^(3)^ = −0.0197) spectral mode of *e*(*a*, *b*), mainly localized on hydrophobic amino acids (A to P).

Strikingly, we find that the mean matrix *e* and its top three spectral modes display some physicochemical interactions at the amino-acid scale, consistent with the MJ energy matrix 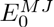, cf. Fig. 2. The first spectral mode (λ^(1)^ = −0.0923) is indeed related to electrostaticity, the second (λ^(2)^ = 0.0363) and third (λ^(3)^ = −0.0197) modes are mainly localized on some hydrophobic amino acids (A to P). The third mode illustrates favorable residue pairing between amino acids of opposing size: A on one hand (Van der Waals volume of 67 Å^3^) and F, I, L on the other hand (Van der Waals volume of 135 Å^3^, 124 Å^3^, and 124Å^3^ respectively). This coevolutionary effect derives from stericity, and is dominant here because of the abundance of the involved amino acids. The favorable interaction between amino acids of opposite size, and unfavorable between amino acids of the same size can be easily understood: given a contact between two amino acids of opposite size, each single change of a small into a large or a large into a small amino acid induces unfavorable steric effects. A compensatory mutation of the second amino acid would be possible.

The sign of all eigenvalues is consistent with what has been previously reported for the MJ energy matrix, cf. Sec. IIE: it is positive for attractive interaction between like amino acids (second mode related to hydrophobicity), negative for attractive interaction between unlike amino acids (first and third modes related to charge and size). Note that the entries and the eigenvalues of *e*(*a*, *b*) are small compared to their counterparts in MJ, a fact we will discuss in Section IIIE.

We conclude that the inferred DCA coupling matrices display quantitative and biologically relevant information, beyond their known efficiency to predict tertiary contacts. However, contrary to the MJ statistical potential (Fig. 1) which includes the possibility of contacts between hydrophilic amino acids (from H to G) and Cysteine-Cysteine (C-C entry) we do not observe such a signal in the modes of the mean matrix *e*. The Pearson correlation coefficient between *e*(*a*, *b*) and 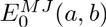 is 0.58.

### C. The C-C signal can be found through structural classification of the pool of Pfam families

The absence of the Cysteine-Cysteine signal may very well be explained by the scarcity of contacts of this type. In order to gain a more detailed view of the possible contact matrices, we divide up the pool of Pfam families into structural domains based on similarities of their structures using the manual Structural Classification of Proteins (SCOP) database^24^ (the repartition is in the Supplementary section V). Five SCOP classes are considered in this analysis : all *α*-proteins, all *β*-proteins, *α*- and *β*-proteins (mainly antiparallel beta sheets: beta-alpha-beta units and segregated alpha and beta regions), membrane and cell surface proteins and peptides, small proteins. The latter is characterized by the abundance of disulfide bridges between two Cysteines. This gives rise to 5 new selections 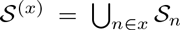, where *x* is the SCOP class (*x* ∈ {*α*,*β*,*α* + *β*, *membrane, small*}). We get 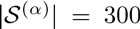, 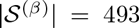, 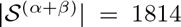, 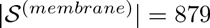, and 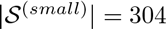.

**FIG. 3:**
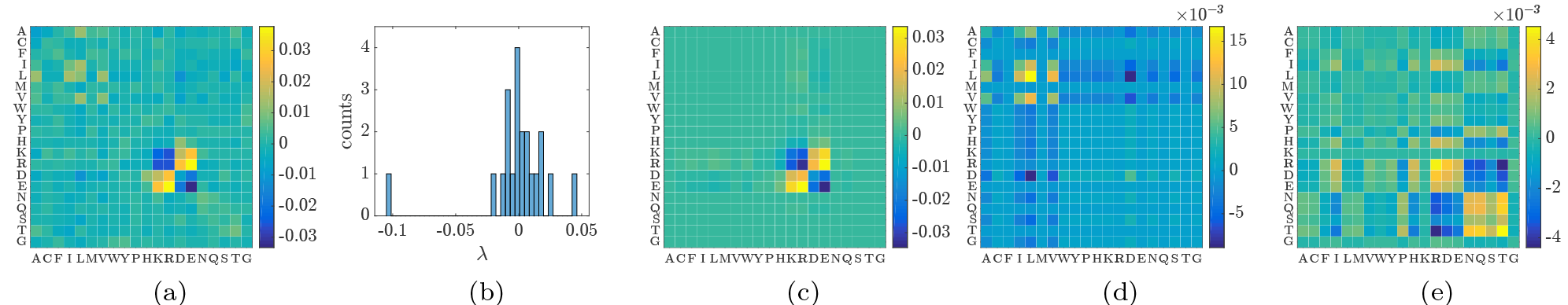
*α* proteins. - (a) **e**(*a*,*b*|*α*) - (b) Spectrum - (c), (d), (e) Top three spectral modes displaying electrostatic (λ^(1)^ = −0.1043), hydrophobic (λ^(2)^ = 0.0459), and hydrophilic (λ^(3)^ = 0.0238) interactions.

**FIG. 4:**
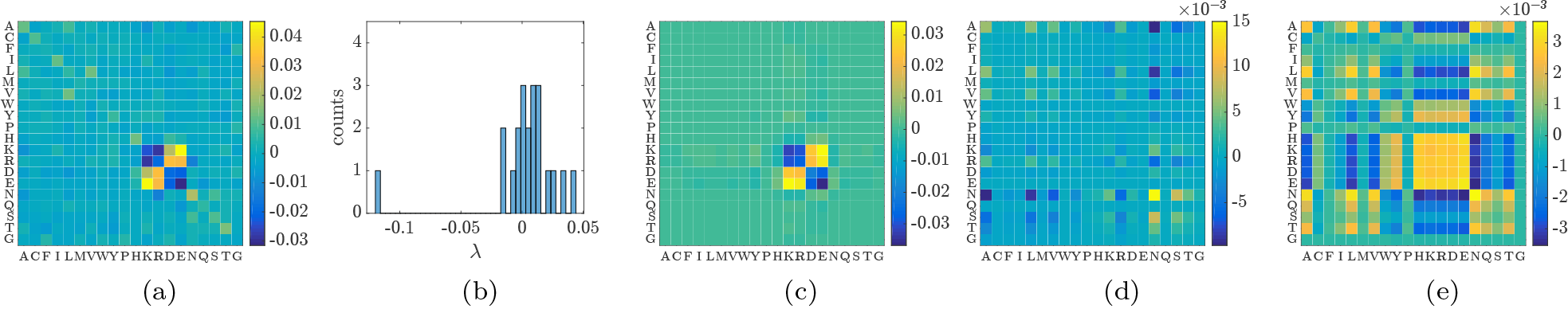
*β* proteins. (a) **e**(*a*,*b*|*β*) - (b) Spectrum - (c), (d), (e) Top three spectral modes displaying electrostatic (λ^(1)^ = −0.1171) and hydrophobic/hydrophilic interactions (λ^(2)^ = 0.0405, λ^(3)^ = 0.0328).

**FIG. 5:**
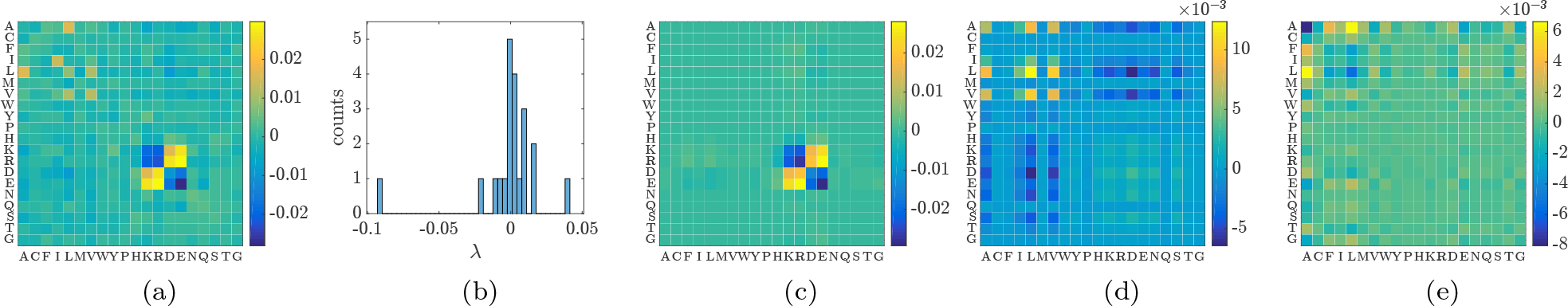
*α* + *β* proteins. (a) **e**(*a*,*b*|*α* + *β*) - (b) Spectrum - (c), (d), (e) Top three spectral modes displaying electrostatic (λ^(1)^ = −0.0905) and hydrophobic (λ^(2)^ = 0.0412, λ^(3)^ = −0.0198) interactions.

**FIG. 6:**
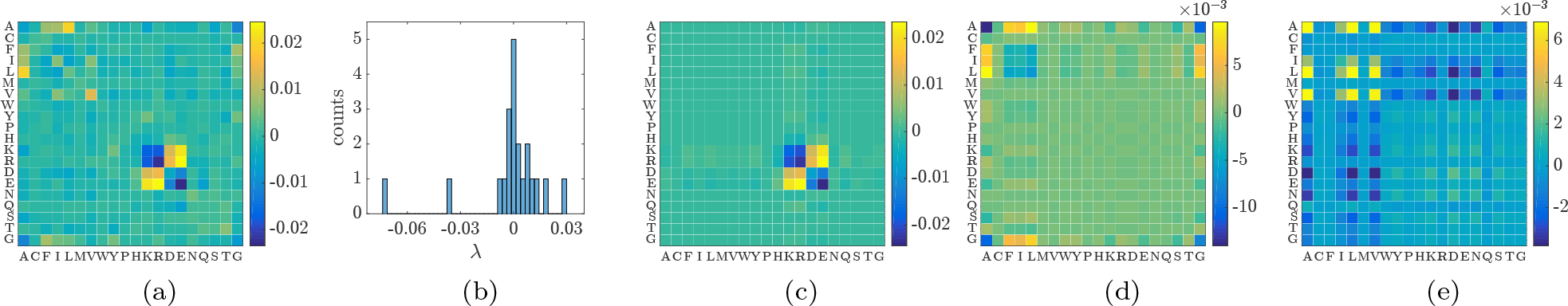
membrane proteins. (a) **e**(*a*,*b*|*membrane*) - (b) Spectrum - (c), (d), (e) Top three spectral modes displaying electrostatic (λ^(1)^ = −0.0729) and hydrophobic (λ^(2)^ =-0.0366, λ^(3)^ = 0.0299) interactions.

**FIG. 7:**
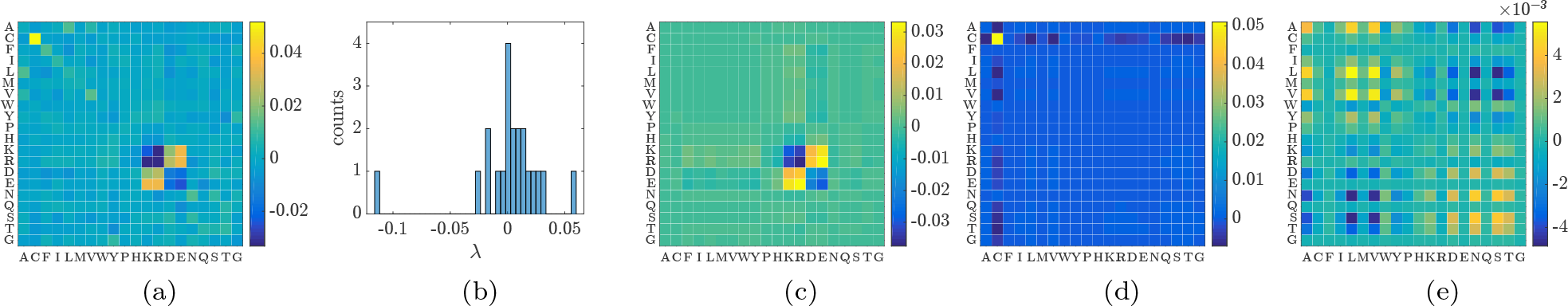
small proteins. (a) **e**(*a*,*b*|*small*) - (b) Spectrum - (c), (d), (e) Top three spectral modes displaying electrostatic (λ^(1)^ = −0.1129, Cysteine-Cysteine (λ^(2)^ = 0.00567), and hydrophobic/hydrophilic (λ^(3)^ = 0.0306) interactions.

Figures 3 to 7 display, for each of the 5 SCOP classes, the new mean matrices**e**(*a*,*b*|*x*) = 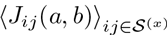, their spectra and the top three spectral modes. Electrostatic spectral modes are found in all 5 SCOP classes (with negative eigenvalues), whereas hydrophobicity-related modes are identified in all but the *small* protein classes. The Cysteine-Cysteine mode is found only in the *small* protein class, as expected (and with a positive eigenvalue). Interestingly, while the hydrophilic signal (amino acids H to G) is still rare in the dominating spectral modes, its presence can be observed in classes *α, β* and *small*, respectively on the third (Fig. 3, panel (e)), second (Fig. 4, panel (d)), and third (Fig. 7, panel (e)) spectral modes. The third mode of *small* (Fig. 7, panel (e)) even displays both hydrophobic *and* hydrophilic interactions, similarly to the MJ energy matrix 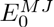 (see Fig. 1, panel (c)).

The spectrum of *e*(*a, b|β*) is dominated by one eigenvalue (λ^(1)^ = −0.1171), the second and third eigenvalues being relatively close (λ^(2)^ = 0.0405, λ^(3)^ = 0.0328). It causes the separation between the second and third spectral modes (Fig. 4, panels (d) and (e)) to be less clear and more sensitive to finite sampling noise than for the other classes, whose spectra are dominated by more than one eigenvalue.

### D. Hydrophilic contacts can be identified considering solvent exposure

The weakness of a signal involving hydrophilic amino acids (from H to G) may be explained by the scarcity of contacts between two sites localized on the surface of the protein as compared to all other contacts-surface amino acids are indeed most likely to be hydrophilic. We now divide the selected residue pairs in *S* into 3 classes depending on the solvent exposure-measured by the relative solvent accessibility (RSA) determined using the naccess software^25^ - of the involved residues, regardless of the Pfam family they are issued from:

- “surface-surface” contacts: more than half of the surface of both residues is exposed to the solvent (selection 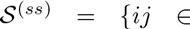 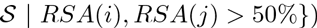,
- “core-core” contacts: less than half of the surface is exposed 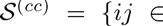 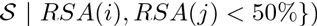,
- “core-surface” contacts: one residue has more than half of its surface exposed, the other has less than half (selection 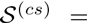 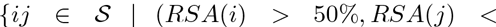 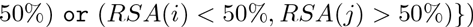.

Fig. 8 displays the repartition of core-core (blue), surface-surface (green), and core-surface (yellow) contacts among all existing tertiary contacts (left panel) and contacts in the selection 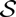 (right panel). As expected, by far the largest part of the tertiary contacts lies in the core of the proteins. Only 2-3% of the (selected) contacts are between surface residues.

**FIG. 8:**
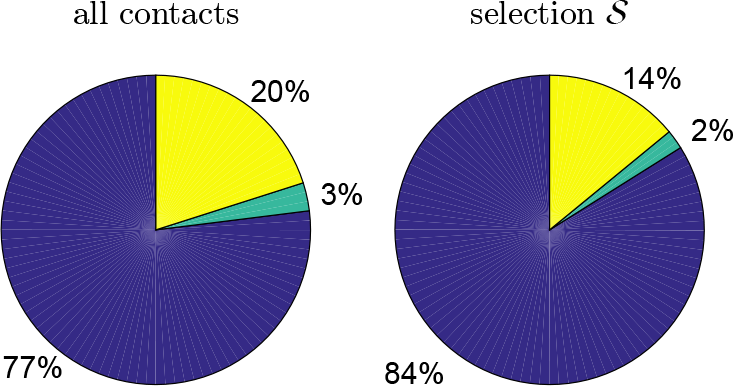
Distribution of core-core (blue), surface-surface (green), and core-surface (yellow) contacts among all contacts (left panel) and contacts in our selection (right panel). Surface-surface contacts are statistically underrepresented in both cases.

**FIG. 9:**
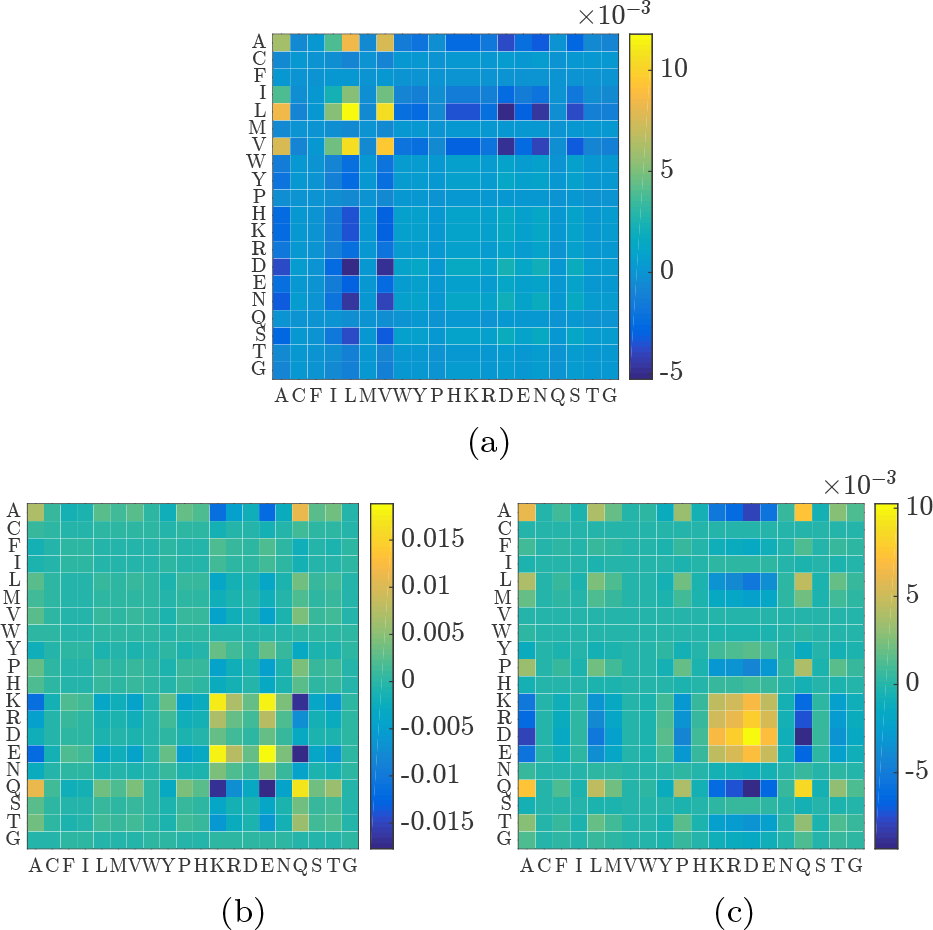
Second spectral modes of the mean matrices (a) *e*(*a, b|cc*) over “core-core” contacts, (b) *e*(*a, b|ss*) over “surface-surface” contacts, and (c) *e*(*a, b*|*cs*) over “core-surface” contacts. A hydrophilicity-related signal is displayed on the 2 latter.

Similarly to what has been done before, we consider average coupling matrices for these 3 new classes: 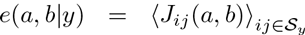, with *y* ∈ {*ss*,*cc*, *cs*} along with their spectral modes. For all classes the first spectral mode displays the usual electrostatic signal, cf. Supplementary section I for a full description of the modes. However, while the second mode of the “core-core” class is localized on hydrophobic amino acids (from A to P) only, in agreement with what is observed on Fig. 2, the second modes of the “surface-surface” and “core-surface” classes are localized only on hydrophilic (H to G) amino acids, as shown on Fig. 9.

### E. Differences with Miyazawa-Jernigan’s statistical potential

The analog of MJ’s contact energy (see Eq. (9a) in^8^) in our description would be approximately the quantity *E*^*stat*^(*a, b*) defined through:

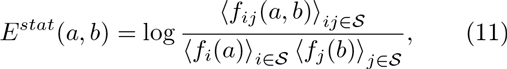

where 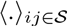 denotes the mean over all residue pairs in the selection 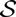 (all Pfam families taken together), and 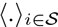 and 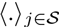 are the means over all single residues involved in a contact pair in the selection 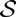. *E*^*stat*^ is then symmetrized and shifted to the zero-sum gauge, cf. Eq. (5). Its first spectral modes are very similar to the genuine MJ energy matrix 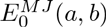( and the Pearson correlation coefficient between E^stat^ and 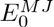 is 0.81 (see Supplementary section II) . Note that, in the zero-sum gauge, the denominator of Eq. (11) is irrelevant.

The *E*^*stat*^ matrix can be related to the inferred couplings in an approximate way as follows. For pairs of site *i*, *j* in contact (in the selection 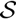), contrary to sites not in contact, the major contribution to the direct coupling *J*_*ij*_ (*a*, *b*) comes from the direct correlation *f*_*ij*_(*a*,*b*)/(*f*_*i*_(*a*)*f_*j*_*(*b*)) between the sites. Indirect contributions to *f*_*ij*_(*a*, *b*), mediated through other sites, are expected to be much smaller. Approximating *J*_*ij*_(*a*, *b*) with log(*f*_*ij*_(*a*,*b*)/(*f*_*i*_(*a*)*f_*j*_*(*b*))) is indeed exact in the case of two interacting sites only. Consequently we introduce the matrix *E*^*DIR*^(*a*, *b*) as

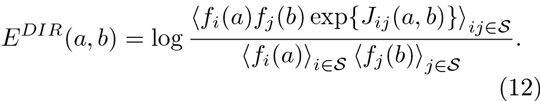

Again, *E*^*DIR*^ is symmetrized and shifted to zero-sum gauge. As displayed on Fig. 10, the first spectral modes are very close to the MJ energy matrix (Fig. 1), although not in the same order (of decreasing eigenvalue in absolute value). The order of magnitude of *E*^*DIR*^ (*a*, *b*) and its top eigenvalues are much more similar to the MJ matrix 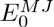, with a Pearson correlation coefficient of 0.77.

**FIG. 10:**
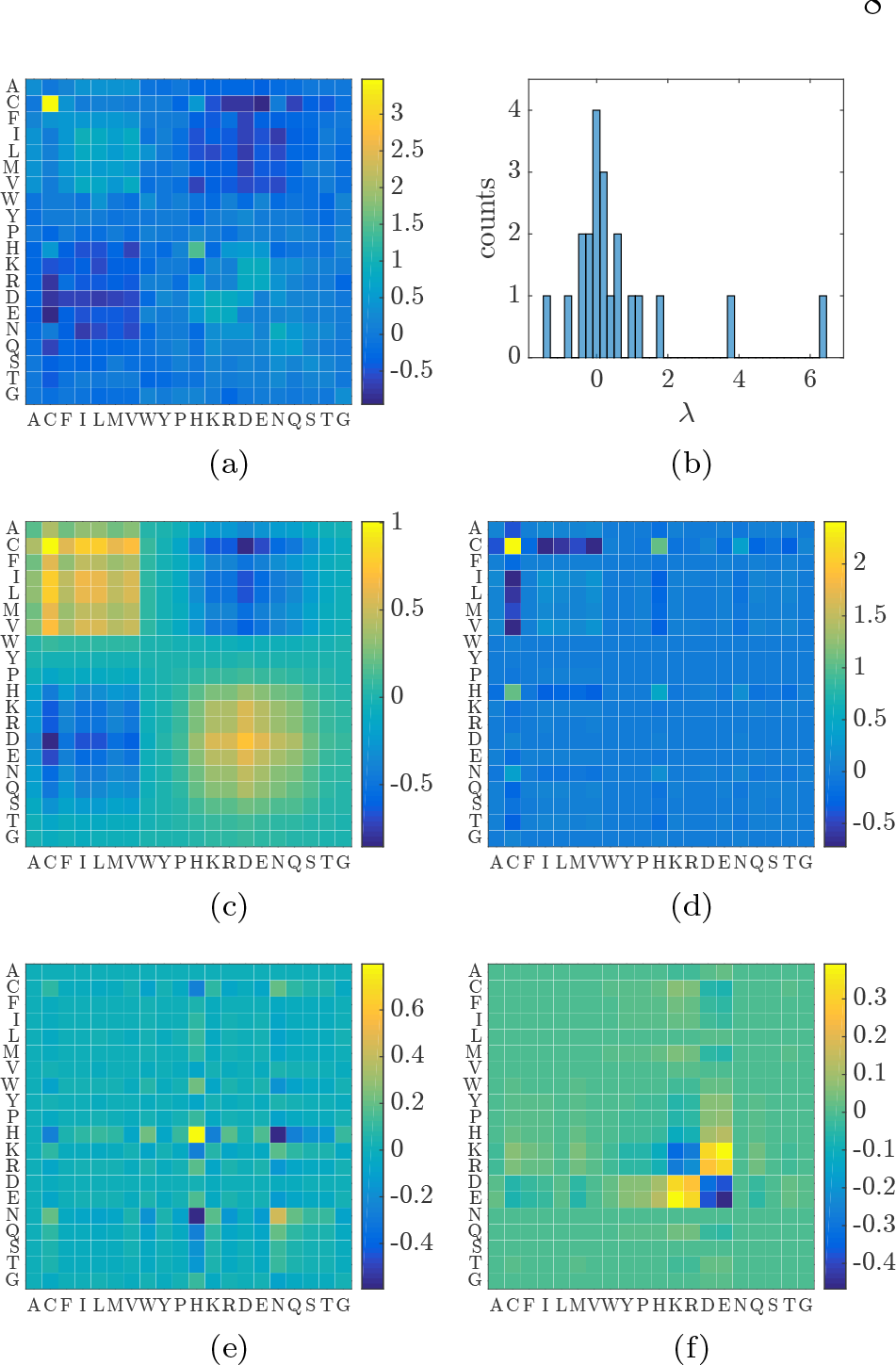
(a) mean matrix *E*^*DIR*^ over all residue pairs in the selection, taking all Pfam families together. (b) Histogram of the spectrum of *E*^*DIR*^. (c), (d), (e), (f) First spectral modes of *E*^*DIR*^ displaying hydrophobic-hydrophilic (λ^(1)^ = 6.44), Cysteine-Cysteine (λ^(2)^ = 3.78), Histidine-Histidine (λ^(3)^ = 1.80), and electrostatic (λ^(4)^ = −1.41) interactions.

This shows that the DCA couplings reflect the full information of the MJ contact energy, provided that the mean is properly weighted by the single sites frequencies. This is consistent with the previous results where the data set of coupling matrices is divided up into structural classes or solvent exposure related classes.

### F. Distance distribution

Within the SCOP classification defined in section IIIC, we assign each residue pair (*i*, *j*) in the selection 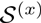to one spectral mode (*k*) of *e*(*a*, *b*|*x*) (with *x* ∈ {*α, β, α* + *β*, *membrane*, *small*}) as follows. We first define the score 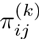 *via* the projection of the coupling matrix *J*_*ij*_ (*a*, *b*) onto the spectral mode (*k*) through

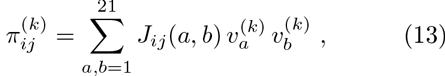

where the 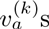 are the components of the eigenvector associated to the *k*^*th*^ eigenvalue of *e*(*a*, *b*|*x*). Then, the residue pair (*i*,*j*) is assigned to the mode (*k*) on which the projection 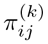 is maximum.

**FIG. 11:**
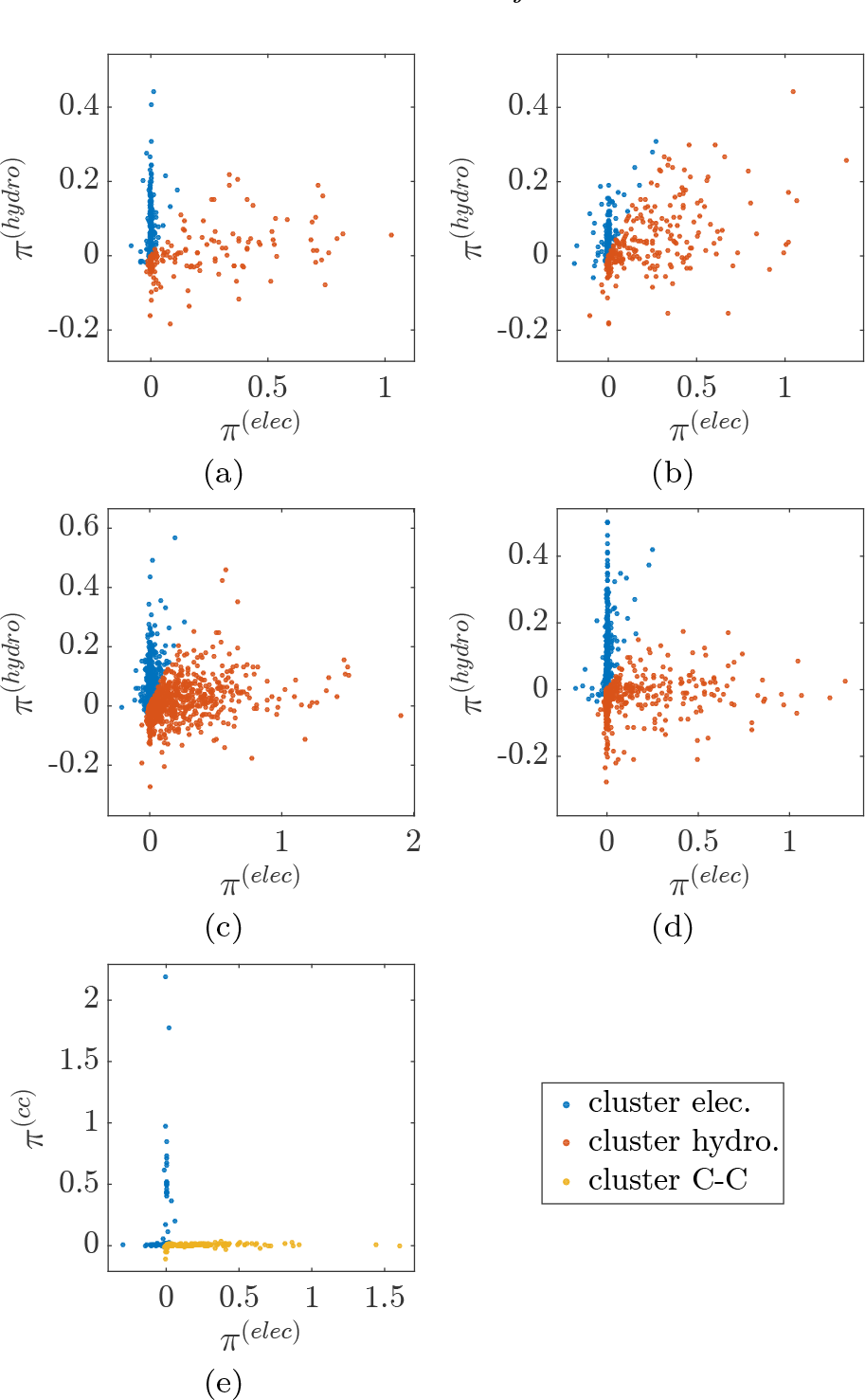
Projection scores 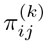, *k* = 1,2 for all residue pairs (*i*, *j*) within SCOP classes (a) *α* (electrostatic and hydrophobic), (b) *β* (electrostatic and hydrophobic), (c) *α*+*β* (electrostatic and hydrophobic), (d) membrane (electrostatic and hydrophobic), and (e) small (electrostatic and Cysteine-Cysteine). Colors indicate the cluster the residue pair has been assigned to: electrostatic (blue), hydrophobic (red), and Cysteine-Cysteine (yellow).

For each class SCOP, we consider the projection onto the top two spectral modes *k* = 1, 2: electrostatic and hydrophobic for the SCOP classes *α, β, α* + *β*, *membrane*, and electrostatic and Cysteine-Cysteine for the class of *small* proteins (Figs. 3 to 7). The top two eigenvalues of *e*(*a, b*|*x*) accounts in each class for about 50% of the sum of all eigenvalues. Figure 11 displays the two projection scores 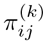, with *k* = 1, 2, for all residue pairs (*i*, *j*) within the five SCOP classes. Each color corresponds to the cluster the residue pairs are assigned to, *i.e.* the mode (*k*) with maximum projection 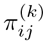.

The projection 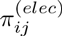 on the electrostatic modes (red dots on Fig. 11) is positive for the vast majority of contacts (*i*, *j*), reflecting the strength and importance of the electrostatic interaction. Residue pairs assigned to hydrophobic modes (blue dots on Fig. 11) usually have a projection *π*^(*elec*)^ close to zero, reflecting the fact that hydrophobic residues are uncharged. While the assignment procedure seems to be well justified for the SCOP classes *α*, *membrane*, and *small* (panels (a), (d), (e)), no clear separation is observed for classes *β* and *α* + *β* (panels (b) and (c)), in which the values of the projection scores of contacts (*i*, *j*) may be both large and comparable in magnitude. This can be explained by the overlapping supports of the electrostatic and hydrophobic spectral modes in theses classes, the latter also having a hydrophilic signal (amino acids K,H,R,D,E are charged *and* hydrophilic), especially for the *β* class, see Fig. 4 panel (d) and Fig. 5 panel (d). Notice that, for the class *small*, the separation between electrostatic and Cysteine-Cysteine modes is very good as the amino acids supporting those interactions are disjoint (K,H,R,D,E for the former, C for the latter).

**FIG. 12:**
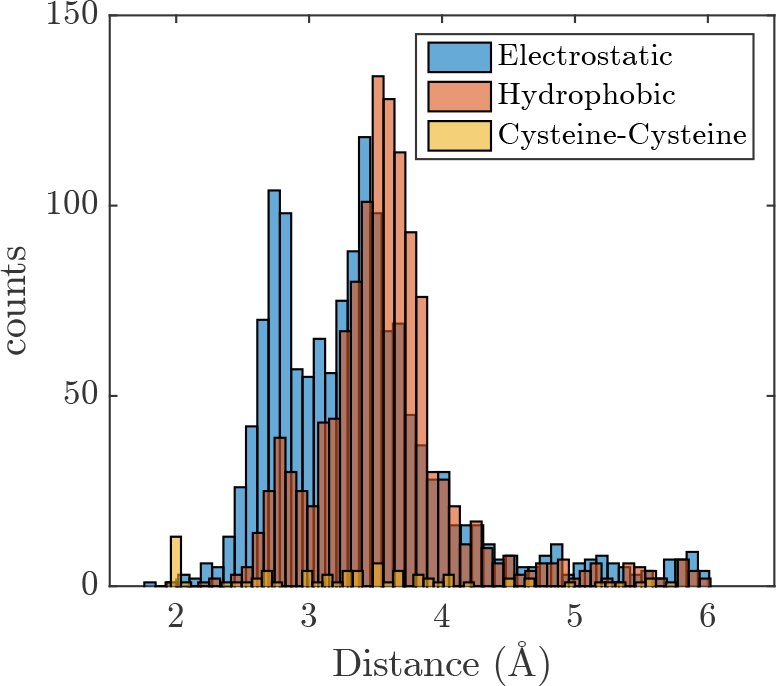
Distribution of distances among the selected residue pairs in contact for the different interaction types, pooled across the SCOP classes.

We now study how the native distances in the tertiary structure between the residue pairs vary with the type of interactions they have been assigned to (electrostatic, hydrophobic or Cysteine-Cysteine) as described above. The distance distributions are shown on Fig. 12, and vary considerably with the interaction types. The “hydrophobic” type involve residue pairs with a contact distance centered around 3.5 Å, the “electrostatic” type displays a bimodal distance distribution mostly around 2.7 Å and 3.5 Å, and the “Cysteine-Cysteine” type is the only one to have a significant number of pairs in contact at short distance 2 Å. Notice that 3.5 Å is the typical distance between heavy atoms, twice the Van der Waals distance (1.7 Å), 2.7 Å corresponds to the distance between atoms linked by a strong to moderate hydrogen bond^26^, and 2 Å is the distance between two Cysteine involved in a disulfide bridge.

## IV. Lattice proteins

Lattice proteins (LP) are exactly solvable models of proteins, folding on a 3D lattice into a compact conformation given by a self-avoiding walk on a cube of dimension 3 × 3 × 3^9^. Real proteins and LP share many common properties (efficient folding, non trivial statistical features, existence of families in the profile HMM sense with conserved folds, etc.), but LP as *in silico* systems allow for precise numerical control. It is easy to generate even large samples of sequences (MSA) corresponding to a single fold, defining the equivalent of a protein family, without any phylogenetic sampling bias. LP are therefore an ideal benchmark for studying and better understanding inference methods developed in the context of real protein data^10^. We will hereafter use the LP framework to study in detail the effect of sampling quality *υs*. regularization strength in the inference of the coevolutionary couplings *J*_*ij*_ (*a*, *b*).

### A. Background

A lattice protein is a chain of *L* = 27 residues occupying the sites of a 3 × 3 × 3 simple cubic lattice; each residue position in the chain can be occupied by one of the 20 different amino acids. 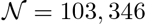 self-avoiding conformations unrelated through symmetry have been enumerated^9^. Each conformation defines a possible structure, or fold of a the protein sequence. The geometry of the cube imposes exactly 28 contacts (neighbors on the lattice but not on the backbone) between the protein sites, cf. Fig. 13.

Given a fold *S*, an energy is assigned to each amino-acid sequence ***A*** = (*a*_*1*_, …,*a*_27_)

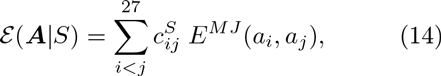

where 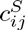 is the contact map of structure *S*, *i.e.* the 27 × 27 adjacency matrix (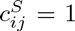 if *i* and *j* are in tertiary contact - not along the chain - and 0 otherwise). Amino acids in contact interact through the MJ statistical potential *E*^*M J*^(*a*,*b*). The probability that a given sequence ***A*** folds in structure *S* is defined by

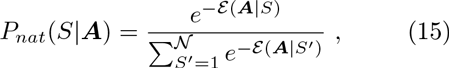

and depends on its energies in all folds *S'*. A good folder is a sequence with a large gap between its energy in the native structure *S* and all the other folds *S'*.

**FIG. 13:**
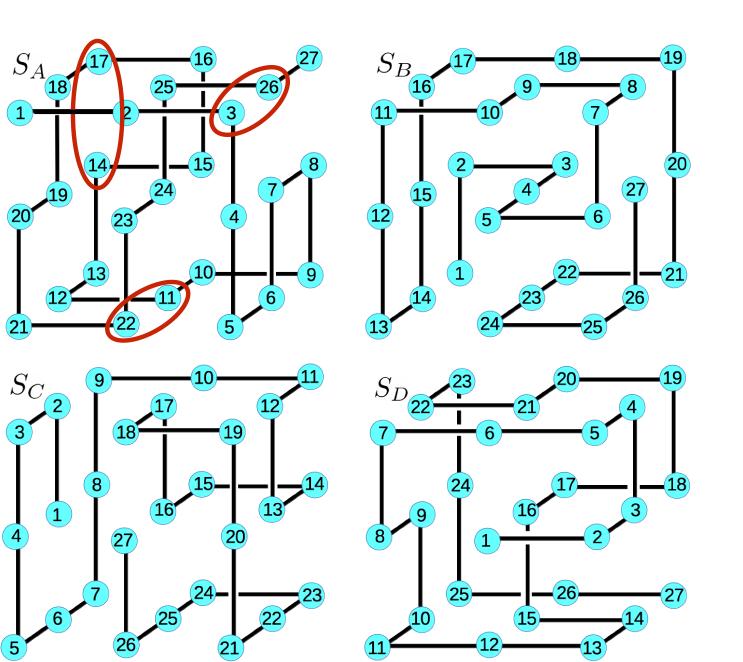
Four representative LP structures used for the analysis. Three among the 28 contacts of structure *SA* have been circled in the top left panel.

Covariation properties of LP were recently studied by Jacquin et al.^10^. MSAs corresponding to the four folds *S*_*A*_, *S*_*B*_, *S*_*C*_, *S*_*D*_ on Fig. 13 were generated by Monte Carlo Markov Chain (MCMC) sampling of *P*_*nat*_(*S*, **A**). The same inverse methods based on Maximum-Entropy and Potts modeling used for real proteins (mean field, plmDCA and the Adaptive Cluster Expansion of^27,28^) were applied to infer pairwise couplings *J*_*ij*_(*a*, *b*) from the one-and two-point statistical correlations measured on the MSAs of the lattice proteins. As in real data, inferred couplings are excellent predictors of contacts in the structure. Interestingly, a linear dependency was observed between the inferred couplings *J*_*ij*_(*a*, *b*) and and MJ energetics parameters *E*^*M J*^(*a*, *b*) used to compute the energy (see Eq. (14)), both in the zero-sum gauge and for a given residue pair (*i*, *j*): 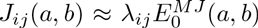. The prefactor λ_*ij*_ was interpreted as a measure of the coevolutionary pressure on the residues (*i*, *j*), due to the design of the native structure. Large positive λ_*ij*_ indicate positive design, and generally correspond to residues (*i*, *j*) in contact in the native structure, but not in its competitor folds *S'*. Conversely, large negative λ_ij_ reflect negative design and generally correspond to residues (*i*, *j*) in contact in competitor structures but not in the native structure^10^. Notice that a profile-HMM^29,30^built on a subpart of a MSA associated to a given fold is very family-specific, and gives high scores to sequences with a high *P*_*nat*_ for this fold. Sequences belonging to other families have lower scores, see Supplementary section III.

### B. Properties of the inferred couplings

We have downloaded the MSAs for structures *S*_*A*_,*S*_*B*_, *S*_*C*_, *S*_*D*_ from the Supporting Information of^10^; each MSA contains *M* = 50000 sequences folding with probability *P*_*nat*_ > 0.995. For each fold, the coupling matrices are computed using plmDCA in zero-sum gauge (as in section III) for 4 different values of the sampling and regularization parameters:

- large sample size (*M* = 50000 sequences) and strong regularization (*μ* = 10^−2^, standard value for plmDCA),
- large sample size (*M* = 50000 sequences) and weak regularization (*μ* = 1/*M* = 2 × 10^−5^),
- small sample size (*M* = 500 sequences extracted from the MSA) and strong regularization (*μ* = 10^−2^),
- small sample size (*M* = 500 sequences extracted from the MSA) and weak regularization (*μ* = 10^−4^).

As expected, the inferred coupling matrices are closely related to the MJ energy matrix^10^, but varying the sampling and regularization strength provide interesting insights. The default regularization parameter is set in plmDCA to the value *μ* = 10^−2^ giving the best results for contact prediction^12^. This regularization strength penalize large couplings and sparsifies the 20 × 20 matrix. With smaller regularization penalties *μ* = 10^−5^ – 10^−4^, couplings can acquire larger values.

**FIG. 14:**
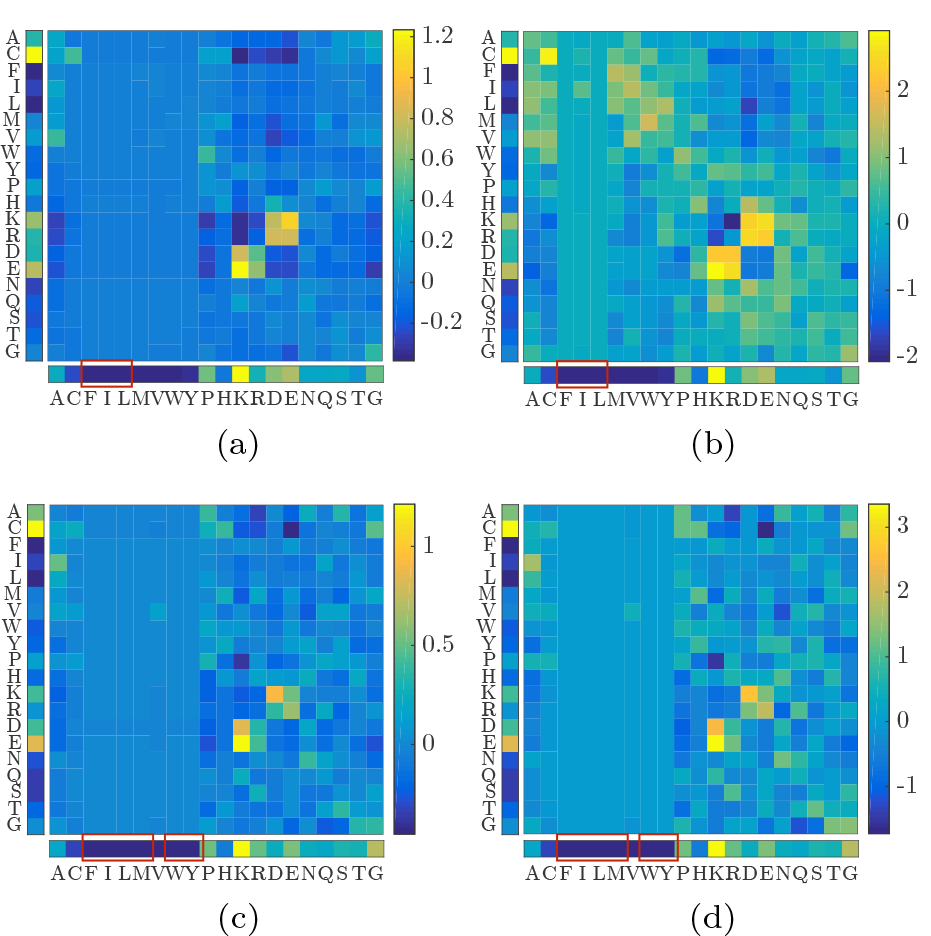
Coupling matrices of pair (14,17), structure *S*_*A*_. Left and bottom colorbars are single site frequencies *f*_14_ and *f*_17_. Red squares indicate zero frequency. (a) *M* = 50000, *μ* = 10^−2^, (b) *M* = 50000, *μ* = 2 × 10^−5^, (c) *M* = 500, *μ* = 10^−2^, (d) *M* = 500, *μ* = 10^−4^.

#### 1. Effect of the regularization

Figure 14 displays the coupling matrix *J*_14_,_17_ of a representative residue pair (14,17) in contact in structure *S*_*A*_ (Fig. 13) at strong (*μ* = 10^−2^, panel (a)) and weak (*μ* = 1/*M* = 2 × 10^−5^, panel (b)) regularizations. Left and bottom colorbars are single site frequencies *f*_14_ and *f*_17_, and red squares indicate zero frequency. The characteristics of the mean coupling matrix will be described in section IV B 3.

Strikingly, decreasing the regularization strength enables new interaction signals to emerge, *e.g.* hydrophobic and Cysteine-Cysteine interaction, which are consistent with the MJ matrix, see panel (a) of Fig. 1. The correlation between *J*_*ij*_ (*a*, *b*) an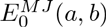 for all (*i*, *j*) in contact in the four studied folds therefore increases, with an average Pearson coefficient raising from 0.51 (strong regularization) to 0.70 (weak regularization).

The unveiling of interactions at weak regularization depends, however, on the amino-acid statistics on the involved sites. For example, for the pair (14,17) displayed on Fig. 14, electrostatic and hydrophilic amino acids (H to G) have sufficiently large frequencies on sites 14 and 17 to produce enough correlation statistics for the corresponding interaction. On the contrary, no interaction signal is revealed at low regularization for amino acids F, I and L, as they are never found on site 17 (vertical band of zero couplings on panel (b)). Decreasing the regularization in the latter case merely results in increasing noise, as discussed in the next subsection.

#### 2. Effect of the sampling

The length of LP is *L* = *27*, which is small compared to real biological proteins (typically 50 – 500 amino acids in a single domain). Moreover, the MCMC procedure used to generate MSAs ensures that the sequences are well distributed in sequence space. In consequence, inference based on good sampling (*M* = 50000 sequences) becomes very accurate. As discussed in Section IIB, the situation for real biological sequences is less optimal. For real biological sequences, the effective number of sequences *M*_*eff*_ is much smaller (we have chosen 500 as a lower bound for the 70 PFMA families studies in the present work), and only very few proteins reach values close to *M* = 50000 chosen for LP in^10^.

To test our analysis of LP in a more realistic situation, we therefore select subalignments of *M* = 500 sequences for each of the four structures. The bottom panels of Fig. 14 display the coupling matrices obtained in this poor sampling situation, at strong (panel (c)) and weak (panel (d)) regularizations. Contrary to the good sampling case, no new interaction signal compatible with MJ is revealed at low regularization. Globally, the coupling matrices of all residue pairs in contacts are even less correlated with MJ, as the Pearson correlation goes from 0.42 (small sample size, strong regularization) down to 0.36 (small sample size, weak regularization). The difference between couplings at strong and weak regularization seems to be due to noise for poor sampling.

The couplings for real protein sequences have been inferred at (plmDCA standard) high regularization (*μ* = 10^−2^). Coherently with what has been described in the last paragraph for LP, and since real biological sequences are not very well sampled (*M*_eff_ ≃ 500 – 1000), decreasing the regularization does not change the mean matrices and their spectral modes; they contain simply more noise.

To sum up the effects of the different parameters (regularization and sampling), Table I gathers the Pearson correlation coefficients between *J*_*ij*_ (*a*, *b*) and 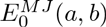 for all amino-acid and residue pairs in the 4 studied folds (4 × 28 = 112 pairs). As we have discussed above, with a good sampling, the correlation between *J*_*ij*_ (*a*, *b*) and 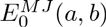 globally increases when the regularization decreases. On the contrary, with poor sampling (as it is the case for real biological data), the correlation slightly decreases when the regularization decreases. However, the inferred signal appears pretty stable at strong regularization, which may be a reason why plmDCA needs this high regularization on real protein data.

**Table I:** Pearson correlation coefficients between *J*_*ij*_ (*a*, *b*) and the MJ energy matrix 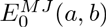 across all residue pairs (contacts / non contacts) in the 4 studied folds for different samplings and regularization strength

| sampling | regularization | correlation |
| --- | --- | --- |
| $M = 50000$ | $\mu = 10^{-2}$ | 0.51 / -0.15 |
| | $\mu = 1/M$ | 0.70 / -0.14 |
| $M' = 500$ | $\mu = 10^{-2}$ | 0.42 / -0.05 |
| | $\mu = 10^{-4}$ | 0.36 / -0.04 |

#### 3. Mean coupling matrix

**FIG. 15:**
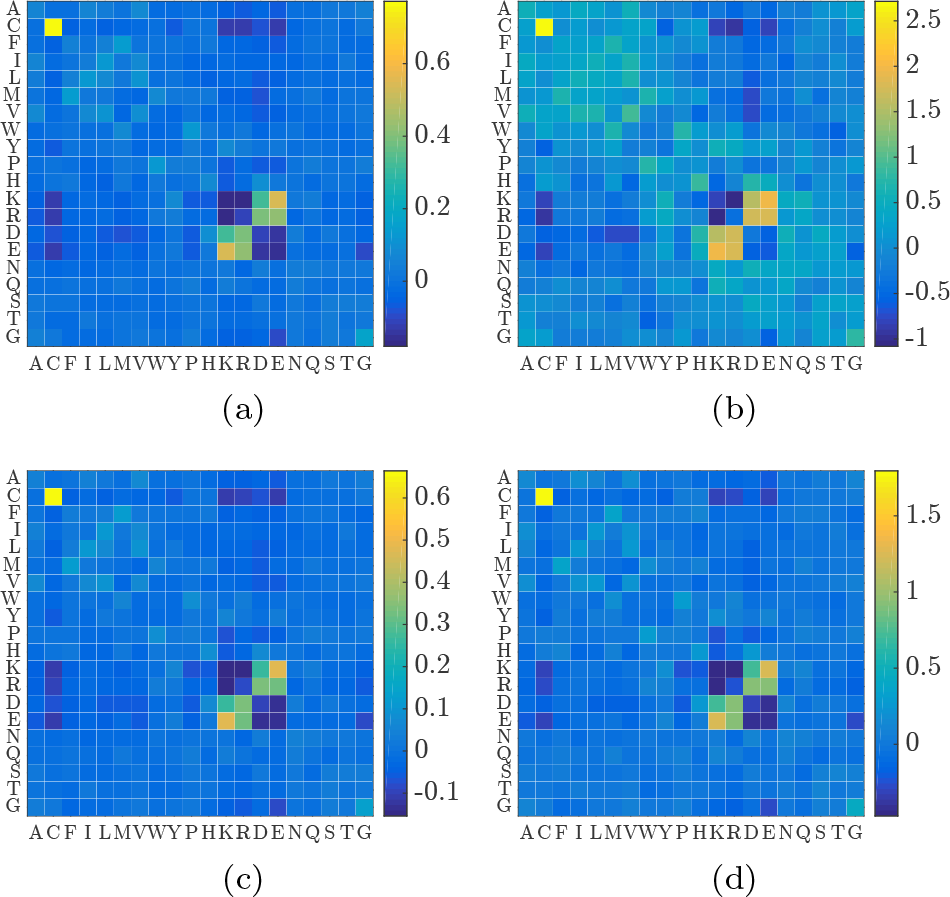
Mean matrix **e**(*a*,*b*|*LP*) over all pairs in contact in the 4 studied folds. (a) *M* = 50000, *μ* = 10^−2^, (b) *M* = 50000, *μ* = 1/*M*, (c) *M* = 500, *μ* = 10^−2^, (d) *M* = 500, *μ* = 10^−4^.

Similarly to what has been done for real sequences data (Section III), we compute the mean matrix

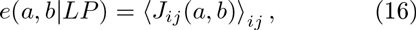

where the mean ⟨.⟩_*ij*_ is over all residues pairs in contact in the 4 studied folds (112 coupling matrices). The 4 cases of different sampling and regularization parameters defined in Section IV B give rise to 4 different matrices *e*(*a*, *b*|*LP*): (*M* = 50000, *μ* = 10^−2^), (*M* = 50000, *μ* = 1/*M*), (*M'*; = 500, *μ* = 10^−2^), and (*M'* = 500, *μ* = 10^−4^), displayed on Fig. 15. Consistently to what has been previously stated, the correlation between *e*(*a*, *b*|*LP*) and the MJ energy matrix 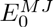 is maximum (0.94) in the case of large sample size and weak regularization (panel (b)). Appendix C reports a full description of the modes of *e*(*a*, *b*|*LP*).

## V. SUMMARY AND CONCLUSIONS

Direct Coupling Analysis exploits the statistical correlations implied by coevolution in protein-multiple sequence alignments to infer residue-residue contact within the tertiary structure. The probabilistic model takes the form of a *q* = 21-states Potts model, whose parameters are inferred to reproduce the one-and two-residue statistics of the data. Usually, the inferred coupling matrices {*J*_*ij*_ (*a*, *b*)} are mapped onto scalar parameters to measure the coupling strength between two residues and thereby predict contacts, without exploring the full information they contain. By studying extensively 70 Pfam protein families, we show that these couplings reflect the physico-chemical properties of amino-acid interactions, such as electrostatic, hydrophobic/hydrophilic, Cysteine-Cysteine and steric interactions. Some of these interaction modes are present in a small fraction of residue pairs only, and are not easily seen in the global analysis over the 70 protein families. We show, however, that Cysteine-Cysteine and hydrophilic signals are unveiled, when we consider the SCOP structural classification (small proteins) and solvent exposure (surface contacts).

Study of lattice proteins (LP) - synthetic protein models folding on a 3D lattice with energetics ruled by the Miyazawa-Jernigan statistical potential - gives useful insights on the effect of regularization strength and sampling on contact classes. Decreasing the regularization penalty (from the default plmDCA value *μ* = 10^−2^ to *μ* = 1/*M*, the inverse of the MSA size) allows for a richer interaction signal to emerge in the coupling matrices, highly correlated with the Miyazawa-Jernigan energy matrix. However, this rich interaction pattern may be inferred only if the sequence sample (MSA) is sufficient large. For sample sizes representative of current real protein databases, decreasing the regularization strength simply makes the correlation with the Miyazawa-Jernigan energy matrix worse as the inferred couplings merely reproduce the sampling noise in the amino-acid pairwise correlations. With such poor sampling strong regularization is more reliable: The inferred interaction signal becomes relatively insensitive to the sample size, explaining why plmDCA on real proteins was found to perform consistently with a constant regularization of *μ* = 0.01. Note that this picture somewhat depends on the inference method considered: more precise inference procedures could allow for detecting a larger correlation with MJ even with poor sampling^11,28,31^.

The order of magnitude of the different mean coupling matrices and their top eigenvalues greatly depend on the regularization strength. Strong regularization imposes important constraints on the couplings, prohibiting large absolute values in the inferred *J_ij_*(*a, b*). On the other hand, LP are characterized by strong structural selection. The presence of negative and positive designs^10^ causes the inferred couplings to be larger. The entries and top eigenvalues of the mean matrices *e*(*a*, *b*|*LP*) are consequently similar or larger than the ones of the MJ energy matrix. The situation for real proteins is less stable, as structure is only partially conserved over protein families, and contacts stabilizing a structure may not always be the same across thousands of distant homologs. This probably explains why the entries and top eigenvalues of the mean coupling matrix *e*(*a*, *b*) are much smaller in real proteins than in the MJ energy matrix.

An important question is whether the detailed structure of the inferred couplings revealed in this work could be used to improve structural predictions, based so far on the Frobenius norms of the couplings only. It was recently shown that for LP the projection of the couplings onto the MJ matrix generally gives a better score for contact prediction than the usual Frobenius-based estimator, see Section IVA and^10^. The reason is two-fold. First the projection, contrary to the norm, has a sign, and allows for the distinction of positive design (positive projection, likely to correspond to contact in the native fold) from negative design (negative projection, likely not to correspond to a contact). Secondly the projection measures the magnitude of the coupling matrix along one direction in the 20 × 20-dimensional space of amino-acid pairs, and is thus not sensitive to the noise in the 399 remaining orthogonal directions, contrary to the Frobenius norm.

However, the applicability to real protein data appears currently limited due to two reasons. First, the projection in^10^ is done on the MJ matrix used in the generative model of the lattice proteins, i.e. complementary information not coming from the data is used. In real proteins, the reference coupling matrix has to be inferred from data first and is thus expected to be less accurate. Second, the currently limited sampling in real proteins was shown to impose a strong regularization during the inference of the DCA model parameters, which even in lattice proteins reduces the correlation between inferred couplings and the MJ matrix. We anticipate this situation to improve soon due to the rapid growth of available genomic data, leading to a better and better sampling of protein families.

Nevertheless, even at the current state of sequence sampling, the coupling matrices contain important quantitative information which can directly be implemented into protein-structure prediction: our work indicates that the type of interaction reflected by the inferred couplings is correlated with the distances in the tertiary structure between the residues in contact (Section IIIF). Cysteine-Cysteine tend to form very strong chemical bonds such as disulfide bridges and therefore are the only contact type associated to very short distances ~ 2 Å. Electrostatic contacts give rise to distances with a bimodal distribution, centered around 2.7 Å and 3.5 Å. Finally, hydrophobic contacts are mainly located around 3.5 Å. While this information has been so far discarded when using DCA or related methods to guide tertiary protein structure prediction, it could in principle be used to make the resulting structural models more accurate.

## ACKNOWLEDGMENTS

We are grateful to H. Jacquin for useful discussions regarding LP, and to E. Westhof and R. Guerois for discussions concerning the structural interpretation of the results. SC, RM and MW are partially funded by ANR-13-BS04-0012-01 (Coevs-tat). AC thanks the Institut des Systèmes Complexes (ISC-PIF) and the Région Ile-de-France for financial support.

## SUPPLEMENTARY

See supplementary material at [URL will be inserted by JCP] for more details on coupling matrices averaged over solvent-exposure related classes (section I), the analog of the Miyazawa-Jernigan matrix computed with one-and two-sites frequencies from alignments (section II), considerations on the profile HMM of lattice proteins (section III), the list of the Pfam families (section IV) and their repartition into SCOP classes (section V), the list of the PDB structures used in the analysis (section VI).

## APPENDIX A: Pseudo-Likelihood Maximization (plmDCA)

The log-likelihood of the data consisting in a MSA of *M* sequences 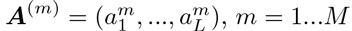, *m* = 1…*M*, reads

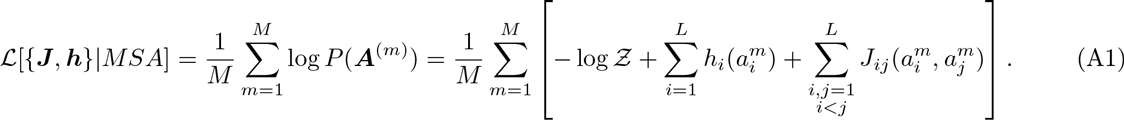

plmDCA substitutes the probability in the log-likelihood in Eq. (A1) by the conditional probability of observing one amino acid at site *r* in a sequence ***A***^(*m*)^ given all the others:

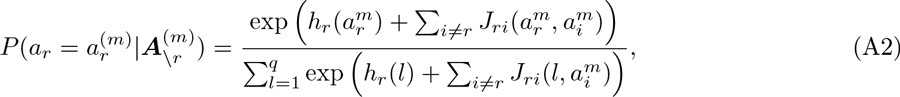

where, for notational convenience, we use *J*_*ri*_ (*l*, *k*)) = *J*_*ir*_ (*k*, *l*) for *i* < *r*. The notation ***A***_*\r*_ =(*a*_1_,…, *a*_*r*−1_, *a*_*r*+1_,…, *a*_*L*_) is used for the subsequence not containing position *r*.

The parameters ***h***_*r*_ and ***J***_*r*_ = {*J*_*ir*_ }_*i*≠*r*_ can be computed via the maximization of the pseudo-loglikelihood

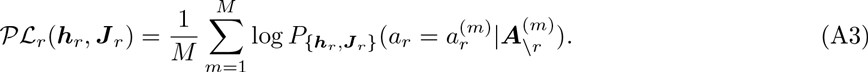

This procedure is statistically consistent, *i.e.* it guarantees to extract the exact parameter values in the limit of an infinitely large sample drawn from the Potts model. However, for a finite sample this procedure returns two different values for the couplings 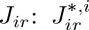 and 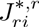 obtained from the maximization of 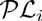 and 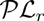 respectively. One simple way to reconcile these values is to replace them by the average: 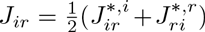. This approach is referred to as *asymmetric pseudolikelihood maximization*^15^, and has been used in this paper.

## APPENDIX B: Crystal structure mapping

We use multiple sequence alignments (MSA) of protein domains downloaded from the Pfam database version 27.0^6^.

We select randomly 70 domain families that satisfied the following criteria: (i) the family contains an effective number of homologous sequences greater than 500, to provide a sufficiently good sample for plmDCA; (ii) each family has at least one member sequence with an experimentally resolved high-resolution crystal structure (resolution lower than 3 Å) available from the Protein Data Bank (PDB)^22^, this permits to extract experimental contact maps and to use the SCOP classification^24^; (iii) every PDB chains that contains a selected domain family has been classified into a unique structural group according to SCOP; (iv) the families are selected to cover a broad range in protein length and to have good sensitivity in the contact prediction.

We consider the first level of SCOP categorization of PDB structures, *the Group*, that account for the types of folds (e.g., beta sheets). 5 structural groups have been used (see the supplementary section V for the list of Pfam families per SCOP class).

A mapping application was developed to map domain family alignments to crystal structures and to extract distances of residue pairs in PDB structures in order to obtain the contact map. Two residues are considered in contact if the minimal distance between all the heavy atoms is lower than 8 Å. This threshold is chosen coherently with prior studies^2^. We take into account several crystal structures, when available, to include the structural variability over homologous proteins that are present in the PDB. Therefore, when more structures are at disposal we take as the distance between residues the minimum distance over the residue pairs in the different PDBs. The complete list of PDB structures can be found in the supplementary section VI.

We compute the relative solvent accessibility (RSA) of a given residue using the naccess tool^25^.

## APPENDIX C: Modes of the mean matrix *e*(*a*,*b*|*LP*), depending on sampling and regularization

*e*(*a*, *b*|*LP*) and its first spectral modes are closest to the ones of the MJ matrix 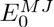 in the case of large sample size and weak regularization (*M* = 50000, *μ* = 1/*M*), as displayed on Fig. 17 and consistently to what has been addressed in section IV B. Table II displays the Pearson correlation coefficients between *e*(*a*,*b*|*LP*) in the 4 cases (panels (a) of Fig. 16 to 19) and the MJ energy matrix 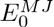.

**Table II:**
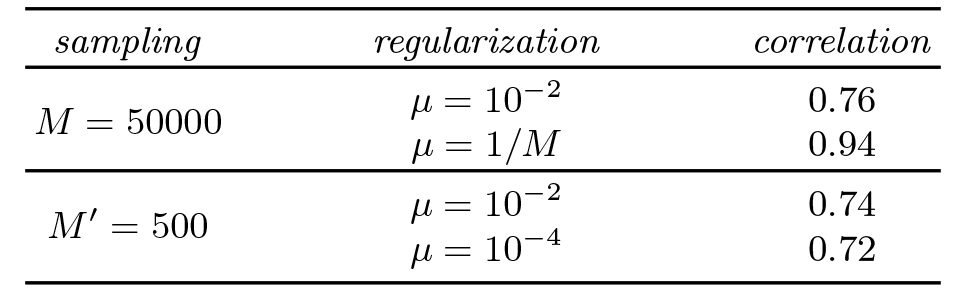
Pearson correlation coefficients between *e*(*a*,*b*|*LP*)) and the MJ energy matrix 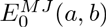 for different samplings and regularization strength

Interestingly, the regularization strength seems to play an important role in determining the order of magnitude of the entries of the matrix *e*(*a*,*b*|*LP*) and its dominant eigenvalues. With a fixed sampling *M* = 50000, the top eigenvalues are divided by 5 with the regularization going from *μ* = 10^−2^ to *μ* = 2 × 10^−5^ (see panels (b) of Fig. 16 and 17). On the contrary, decreasing *M* at fixed regularization does not affect the top eigenvalues (panels (b) of Fig. 16 and 18).

In the optimal case of large sample size and weak regularization, where the correlation with the MJ energy matrix is maximal (see Table II), the entries of *e*(*a*,*b*|*LP*) and its top eigenvalues are larger than the MJ energy matrix (see Fig. 1). Decreasing the folding probability *P*_*nat*_, and therefore the structural constraints, causes the inferred couplings to decrease. It illustrates the strong influence of the evolutionary pressure and positive/negative design in LP^10^.

**FIG. 16:**
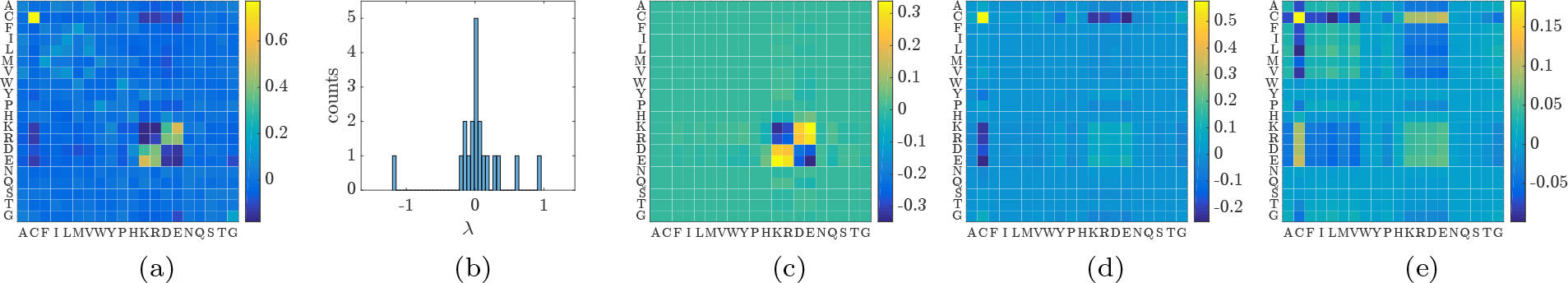
(***M* = 50000**, ***μ* = 10^−2^**). (a) mean matrix *e*(*a*,*b*|*LP*) over all residue pairs in contact across the 4 studied fold. (b) Histogram of the spectrum of *e*(*a*,*b*|*LP*). (c), (d), (e) First spectral modes of *e*(*a*,*b*|*LP*) displaying electrostatic, Cysteine-Cysteine, and mixed Cysteine-Cysteine/hydrophobic/hydrophilic interactions.

**FIG. 17:**
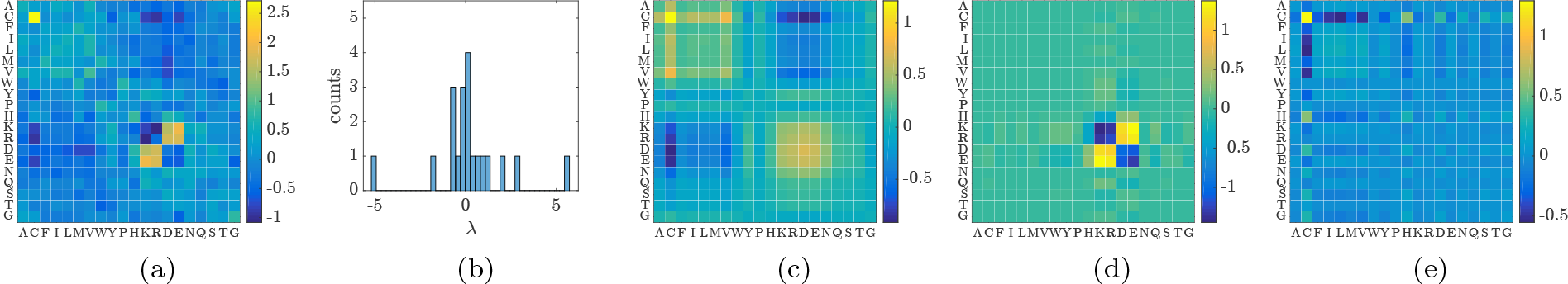
(***M* = 50000**, ***μ*** = **1/*M*** = **2 × 10^−5^**). (a) mean matrix *e*(*a*,*b*|*LP*) over all residue pairs in contact across the 4 studied fold. (b) Histogram of the spectrum of *e*(*a*,*b*|*LP*). (c), (d), (e) First spectral modes of *e*(*a*,*b*|*LP*) displaying electrostatic, Cysteine-Cysteine, and hydrophobic/hydrophilic interactions.

**FIG. 18:**
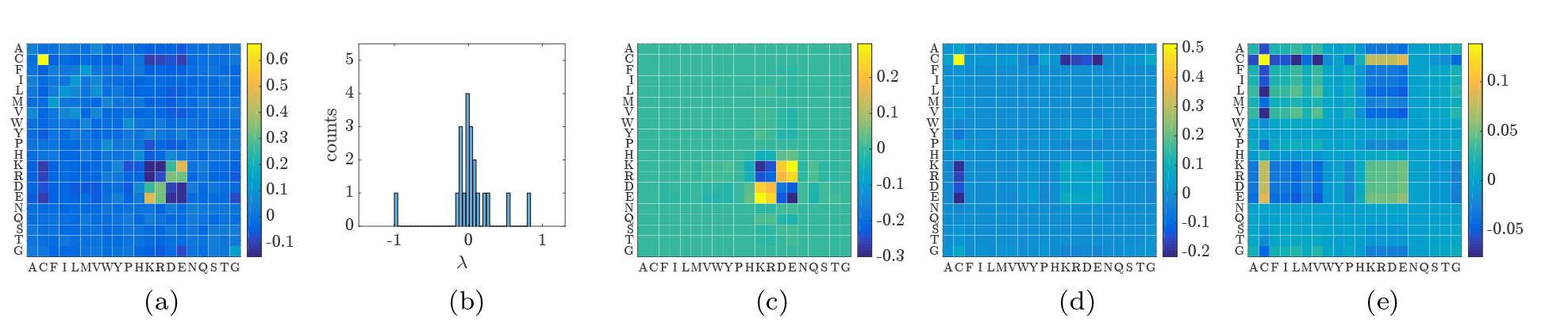
(***M′*** = **500**, ***μ*** = **10^−2^**). (a) mean matrix *e*(*a*,*b*|*LP*) over all residue pairs in contact across the 4 studied fold. (b) Histogram of the spectrum of *e*(*a*,*b*|*LP*). (c), (d), (e) First spectral modes of *e*(*a*,*b*|*LP*) displaying electrostatic, Cysteine-Cysteine, and mixed Cysteine-Cysteine/hydrophobic/hydrophilic interactions.

**FIG. 19:**
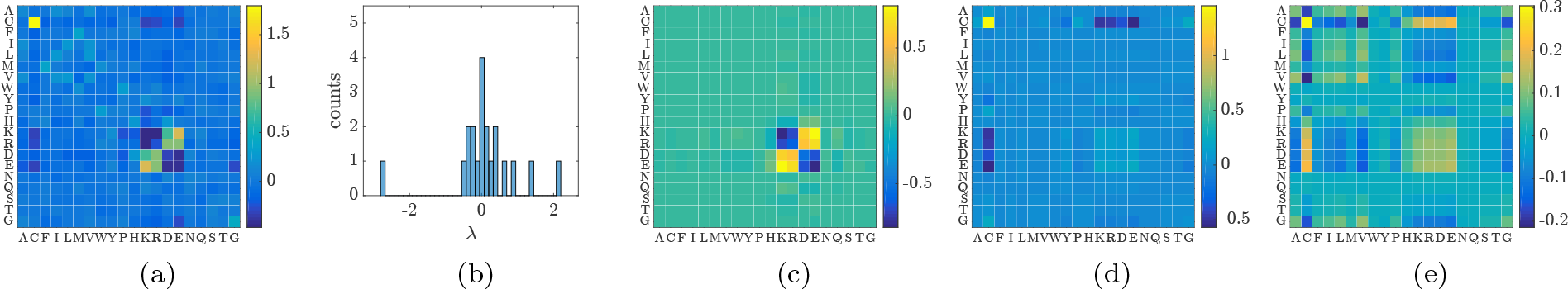
(***M′*** = **500**, ***μ*** = **10^−4^**). (a) mean matrix *e*(*a*,*b*|*LP*) over all residue pairs in contact across the 4 studied fold. (b) Histogram of the spectrum of *e*(*a*,*b*|*LP*). (c), (d), (e) First spectral modes of *e*(*a*,*b*|*LP*) displaying electrostatic, Cysteine-Cysteine, and mixed Cysteine-Cysteine/hydrophobic/hydrophilic interactions.

